# Characterization of Oral Tactile Sensitivity and Chewing Efficiency Across Adulthood

**DOI:** 10.1101/333674

**Authors:** Grace E. Shupe, Zoe N. Resmondo, Curtis R. Luckett

**Author notes:** Corresponding author: Curtis R. Luckett; Department of Food Science, University of Tennessee,2510 River Drive, Knoxville, TN 37996, U.S.

## Abstract

Texture perception is one of the most important factors in food acceptance, yet population-wide differences in texture sensations are not well understood. The variation in texture perception across populations is thought to depend on oral tactile sensitivity and oral processing behaviors. To address this hypothesis, we aimed to measure tactile acuity with a battery of tests and quantitate the relationship to oral processing. The study was performed on 98 participants, in 3 age groups (20-25, 35-45, or over 62). Two main measures of oral sensitivity were performed: to assess bite force, subjects were asked to discriminate between foam samples of varying hardness. Secondly, to assess lingual sensitivity the subjects were asked to identify 3D printed shapes, ranging from 3mm to 8mm, using their tongue, as well as identify confectionary letters. Additionally, chewing efficiency was measured through assessing each participants ability to mix two-colored chewing gum. In general, we found that sensitivity and chewing efficiency in the younger age groups was superior to that of older adults (p<0.0001). We also found a positive linear trend between bite force sensitivity and chewing efficiency with younger participants, a trend not found in older participants. We found no significant relationship between age groups for bite force and chewing efficiency, suggesting that age-related declines in bite force sensitivity are not a significant cause of altered oral processing ability. These results help bolster evidence that sensitivity and oral processing are related, as well previously reported declines in both as people age.

## Introduction

While food texture perception is multisensory in nature, involving sight and hearing, it is mainly routed in touch (Nishinari, Kohyama, Kumagai, Funami, & Bourne, 2013; Szczesniak, 2002). Touch is perceived through pressure, vibration, pain, and stretching (Carlson, 2012). Tactile sensitivity in the mouth, often termed, oral sensitivity, is the ability to determine shape, size, and surface texture of food stuffs (Calhoun, Gibson, Hartley, Minton, & Hokanson, 1992; Engelen, Van der Bilt, & Bosman, 2004). Oral sensitivity has been shown to be dependent on several factors, such as gender, but especially age and dental status (Bangcuyo & Simons, 2017; Calhoun et al., 1992). With age, oral sensitivity decreases along with other physiological measures like fungiform papillae density and dental health (Bangcuyo & Simons, 2017; Calhoun et al., 1992). This loss of sensitivity/oral ability may be a result in discomfort or an inability to adequately prepare a bolus and potentially lead to problems with swallowing; leading to dysphagia in older populations, and therefore a lack of use, resulting in an overall decreased sensitivity (Wada, Kawate, & Mizuma, 2017).

Various methods have been used to determine sensitivity in the oral cavity, these have included oral form recognition (Essick, Chen, & Kelly, 1999), size and weight discrimination tests (Johnson & Phillips, 1981), stereognosis (Jacobs, Serhal, & van Steenberghe, 1998), two-point discrimination (Engelen et al., 2004), force perception (Pigg, Baad-Hansen, Svensson, Drangsholt, & List, 2010), and other physiological measures (Bangcuyo & Simons, 2017; Calhoun et al., 1992; Linne, 2017).

While there is no shortage of methods to assess oral tactile sensitivity, few studies have directly attempted to relate oral sensitivity to elements related to food/beverage intake, most importantly texture perception and oral processing. Recently, Linne et al. investigated the relationship of astringency perception and roughness perception, finding that astringency is related to oral roughness sensitivity for some compounds, but not others (Linne et al. 2017). Additionally, Engelen et al. found the ability of individuals to discriminate sizes of steel spheres to correlate to their masticatory performance (2004). However, performance on a two-point discrimination task was not correlated to masticatory performance, suggesting certain forms of oral tactile sensitivity is more important for oral processing than others.

One area that has yet to be explored is the use of a person’s bite as a physiological measure that could be used to characterize masticatory performance. Masticatory performance and bite force sensitivity have been explored separately, but have not been studied together to determine the relationship to one another (Carlsson, 1974). Bite force measurements are often used in dentistry, as force is applied to an object by the teeth, the many nerve endings innervating the periodontal ligament give the ability to distinguish small changes in pressure. In order to determine jaw placement and avoid discomfort while chewing due to an unintended collision of teeth feedback from these nerve endings is utilized during mastication (Desislava & Mariana, 2016).

It has been suggested that methods being used to determine sensitivity, should focus on how texture (shape, force, size, orientation, etc.) is perceived then relayed back into the masticatory feedback loop (Chen, 2014). The sensitivity to bite force would be expected to be closely related to mastication feedback loop. As mentioned earlier, these questions have not been extensively addressed in the oral cavity. However, studies investigating grip force have detailed the extreme precision in which healthy subjects use enough grip to prevent accidental slips, but not induce muscle fatigue or damage to the object (Johansson & Westling, 1984). Interestingly, the application of topical anesthesia significantly reduces the ability of subjects to use precise grip forces, suggesting that tactile sensitivity is key to this skillset (Johansson & Westling, 1984). Translating this to the oral cavity, sensitivity to bite force may be a key factor in explaining the variation in chewing efficiency. Those who do not possess adequate bite force sensitivity are likely to use too much force and fatigue their masticatory muscles or too little force, leading to poorly masticated food.

The purpose of this study is to better understand the relationships between oral physiology and chewing efficiency (a measure of oral processing). More specifically, we look to quantify the relationship of bite force sensitivity, oral stereognosis, and lingual tactile sensitive to masticatory performance (as measured by chewing efficiency). Secondarily, we seek to look for changes in both oral sensitivity and chewing efficiency across the adult lifespan.

## Materials and Methods

### Participants

Ninety-eight participants were recruited for this study. Participants reported a good sense of smell, had no allergies or food restrictions and were not pregnant. Participants were also asked to self-report common dental procedures such as root canals, crowns, partial or full dentures. Participants were grouped by age as either young (20-25, n=34), middle (35-45, n=31), or old (>62, n=33); see Table 1 for participant demographics. All participants signed an informed consent and were compensated for their time. This experiment was conducted according to the Declaration of Helsinki for studies on human subjects and approved by the University of Tennessee IRB review for research involving human subjects (IRB #17-04120-XP). The authors declare that they do not have any conflict of interest.

**Table 1.** Demographics of participants by age group.

| Demographics |  | Age Group |  |  |
| --- | --- | --- | --- | --- |
|  |  | Young | Middle | Older |
| Age | N | 34 | 31 | 33 |
|  | Mean | 22.5 ± 1.6 | 40 ± 3.1 | 73 ± 6.1 |
|  | Max | 25 | 45 | 87 |
|  | Min | 20 | 35 | 62 |
| Dental Status | Mean | 5 ± 0.38 | 5 ± 0.76 | 4 ± 1.35 |
|  | Max | 5 | 5 | 5 |
|  | Min | 3 | 2 | 0 |
| Gender | Female | 64.7% | 58.1% | 54.5% |
|  | Male | 35.3% | 41.9% | 45.5% |
| Ethnicity | White | 76.5% | 93.5% | 100.0% |
|  | African American | 8.8% | 3.3% | - |
|  | Asian/Pacific Islander | 8.8% | 3.3% | - |
|  | Latino | 5.8% | - | - |

\* Mean values have SD as the error term.

### Stimuli

#### Lingual Sensitivity

Based on Essick et al., an applicator and 10 different shape stimuli (of 4 different sizes in both raised and recessed orientations, see Table 2) were used to determine lingual sensitivity (1999). Geometric shapes were chosen as to refrain from assuming that participants have a familiarity with the Latin alphabet. Sizes were optimized by a pilot study to guard against possible ceiling/floor effects. All materials were 3D printed using a uPrint SE Plus® printer (Stratasys, Eden Prairie, MN) (See Figure 1). The ten shapes consisted of a variety of geometric shapes of varying difficulties and were as follows: square, rectangle, triangle, star, hexagon, circle, half circle, diamond, cross, and heart. The longest axis was used to determine the size in millimeters for each stimulus, and across all four sets the orientation of each shape was not altered and appeared exactly as pictures on the provided answer bank in order to prevent confusion.

**Table 2.**
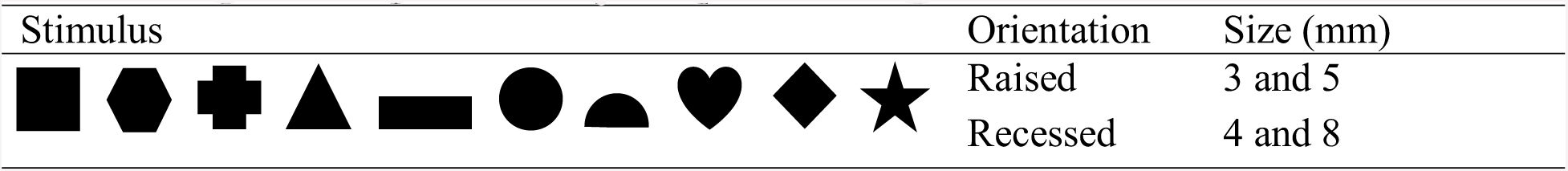
Shape stimuli presented to participants showing all orientation and size combinations.

**Figure 1.**
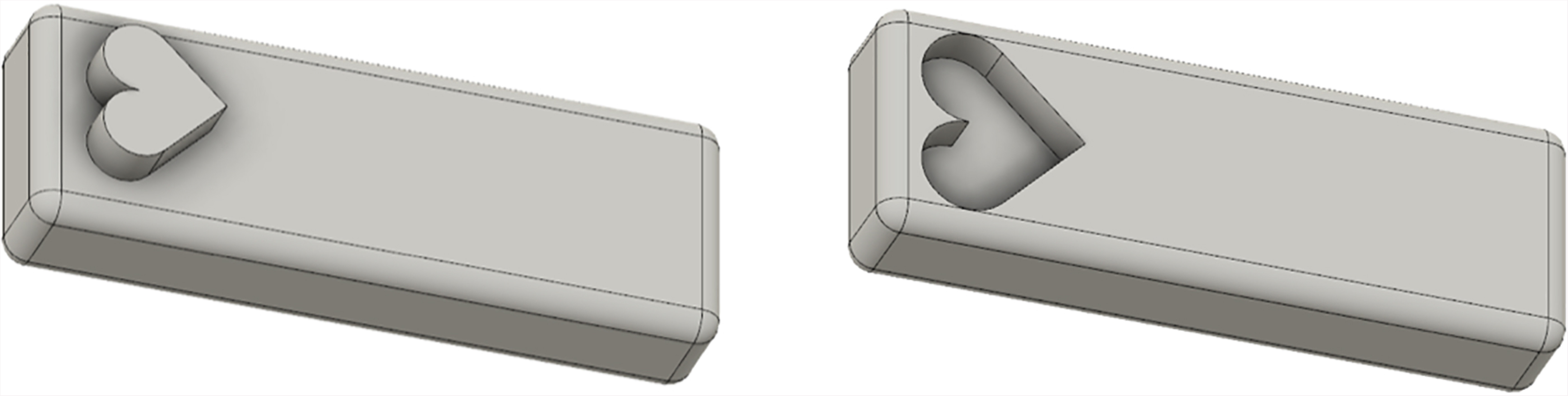
Vector drawing of stimuli: 5 mm raised and 8 mm recessed heart.

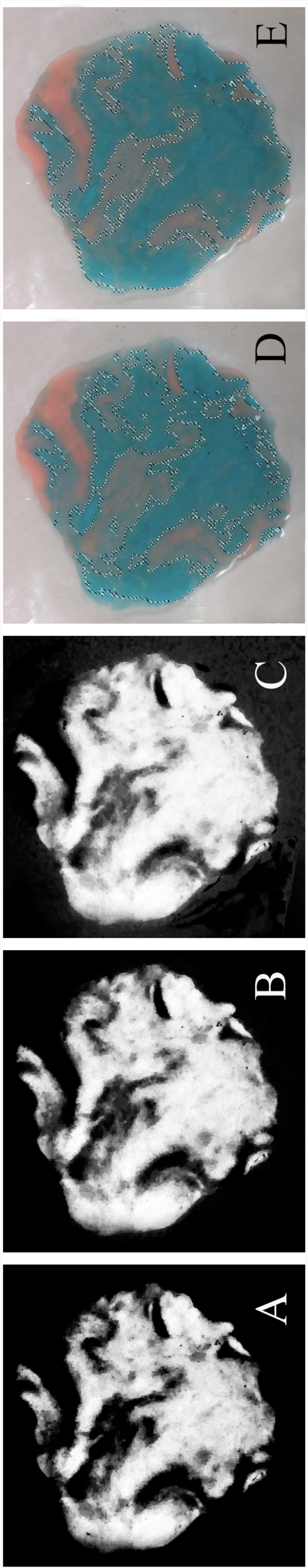

#### Sensitivity to Bite Force

Several foam samples with multiple hardness levels (or compression factors), yet having similar densities were used in this study. Foam was cut into 1 cm cubes and attached to a wooden applicator to allow for the placement of each sample between the molars. Hardness levels were verified using a TA.XT Plus Texture Analyzer and Exponent software (Texture Technologies Corp. and Stable Micro Systems, Ltd., Hamilton, MA) and shown in Table 3.

**Table 3.** Hardness distributions of 1 × 1 cm foam samples using a TA.XT Plus Texture Analyzer.

| Foam # | Mean ± SD (kg) |
| --- | --- |
| 1 | 0.123 ± 0.024 |
| 2 | 0.149 ± 0.009 |
| 3 | 0.191 ± 0.013 |
| 4 | 0.229 ± 0.006 |
| 5 | 0.226 ± 0.033 |

**Table 4.** Pearson’s correlation coefficients (r) among the oral sensitivity and masticatory performance tasks.

|  | Age | Dental Status | Chewing Efficiency | Stereognosis | Lingual Sensitivity | Bite Force Sensitivity |
| --- | --- | --- | --- | --- | --- | --- |
| Age | - | -0.6004** | 0.1755 | -0.4136** | -0.3693* | 0.0005 |
| Dental Status |  | - | 0.2776* | 0.2749* | 0.1665 | -0.1082 |
| Chewing Efficiency |  |  | - | 0.1359 | 0.1199 | -0.0341 |
| Stereognosis |  |  |  | - | 0.4645** | -0.0672 |
| Lingual Sensitivity |  |  |  |  | - | -0.0100 |
| Bite Force Sensitivity |  |  |  |  |  | - |

\* Significant at the 0.01 level.
\*\* Significant at the 0.001 level.

#### Oral Stereognosis

Based on Calhoun et al., confectionary alphabet letters (Haribo® Alphabet Letters Gummy Candy, Haribo of America, Inc., Rosemont, IL) were used to determine stereognosis ability (1992). Letters displaying physical signs of unconformity in letter shape were not used. Using preliminary experimentation, the stimuli were sorted into two groups based on difficulty of recognition. The difficulty of samples was modulated between participants such that each participant received the same ratio of difficult to easily recognized stimuli. Each participant received nine (9) confectionary letters with no letters being repeated.

#### Chewing Efficiency

Using the method defined by Schimmel et al., two different colors (blue and pink) of Hubba Bubba® tape chewing gum (The Wrigley Company Ltd, Plymouth, Devon, England) were used to measure chewing efficiency (2007).

### Procedure

Upon arrival each participant was familiarized with each stimulus and the general tasks to be completed. The presentation of stimuli within each test was randomized, with the overall order of presentation maintained between participants to reduce fatigue. Participants completed one (1) approximately hour-long session with the following serving order; gummy letters (3), shapes, gum, shapes, gummy letters (3), gum, foam, gummy letters (3). Participants were asked to verbally respond with all answers, which were then recorded by members of the research team. Participants also filled out demographics upon completion and were compensated ten dollars for time participating.

#### Oral Stereognosis

Prior to samples being administered participants were instructed that all 26 capital letters of the alphabet were an option, letters were in arial font. Once participants were ready to proceed, they were blindfolded to ensure letters would not be visualized and metal forceps were used to place samples in the mouth. Participants were given as much time as needed to identify the sample, no answer key was given. Once a participant had an answer, they would verbally respond, and answers were recorded by administering personnel.

#### Shape Identification

Participants were familiarized with both orientations (raised and recessed) and shown multiple shapes in different sizes until they were confident in their understanding of the task. Participants were presented with an answer key of all possible shapes. Participants were instructed that each shape would only be used once per size (four sizes), but that they could use the same answer multiple times if desired. Size and order of shapes was randomized, only one size was presented at a time.

#### Chewing Efficiency

Participants were given a gum sample and instructed to chew normally and would be told when to stop and place samples in a plastic bag. Each participant was allowed to chew for 10.0 seconds. We choose not to limit the chewing efficiency measurement by number of chewing cycles due to compensatory strategies exhibited by older adults (Song, Giacalone, Bølling Johansen, Frøst, & Bredie, 2016). Each participant completed this task in duplicate.

#### Sensitivity to Bite Force

A 2-AFC forced-choice paradigm was used to assess sensitivity to pressure. Each 2-AFC consisted of a reference and another sample of varied firmness. All samples were presented in duplicate. Prior to samples being administered participants were familiarized with materials and demonstrated what would be done with a visual by administering personnel. Panelists were asked which side of the jaw they would prefer testing be performed on (the side with the most natural teeth or dominate chewing side). Panelists were then blindfolded and samples places between the back molars of the preferred side monadically with as little time between samples as possible (ensuring that stimuli were correctly oriented and placed between the molars). Participants were allowed to retest if necessary, sample order was maintained.

## Data Analysis

Data was structured as the number of correct responses each panelist gave for all oral sensitivity measures and chewing efficiency was reported as a percentage (averaged over both trials). In order to determine overall oral sensitivity, the sum of correct responses for lingual sensitivity, stereognosis, and bite force sensitivity tasks was used. Dental status was rated on a zero to five scale (5 – filling, 4 – crown, 3 – root canal, 2 – multiple crowns and root canals, 1 – dentures, 0 – minimal natural teeth with no prosthetics), zero and one were considered notably compromised during analysis.

Gum samples were flattened into a 1 mm thick disk, and pictures taken of both sides using an 8.0-megapixel camera (2448×3264). The samples were analyzed using Adobe Photoshop Creative Cloud® (Adobe Systems Inc., San Jose, CA). A reference of un-chewed gum was used the determine the hex code 237a88, which was then used to calculate pixel counts at three fuzziness settings (60, 75, and 90 to account for slight color variation in chewed samples) using the color range selection and the measurement tool (see Figure 2). These measurements were averaged for each side and trial, and the overall pixel count was reported as a percentage of the total number of pixels (chewing efficiency).

All results were analyzed using JMP Pro 13.1 (SAS Institute, Cary, NC), with statistically significant defined as p < 0.05. Differences in lingual sensitivity, stereognosis, bite force sensitivity, and chewing efficiency were examined across age groups by multiple analysis of variances (ANOVAs) and specific LS means contrasts and linear regression was performed for the categorical variable dental status. Pairwise post-hoc comparisons were performed using Tukey’s HSD test and Pearson’s correlations were used to determine associations between measures. To compare oral sensitivity scores, an ANOVA was run, using age group as the sole factor. Each sensitivity task was analyzed separately as well with lingual sensitivity, stereognosis, and bite force sensitivity each compared across the age groups. To compare chewing efficiency ratings, a one-way ANOVA was run, using age and chewing efficiency as a fixed factor.

## Results

### Age

Oral sensitivity was different across the age groups, the older age group having lower total scores than both the young and middle age groups (F_2,95_ = 10.14, p < 0.0001). In looking at specific oral sensitivity measurements, both lingual sensitivity and stereognosis differed across the age groups (F_2,95_ = 7.94, p = 0.0006 and F_2,95_ = 15.02, p < 0.0001, respectively), as shown in Figure 3. Conversely, bite force sensitivity did not differ by age group (F_2,95_ = 0.30, p = 0.739).

**Figure 3.**
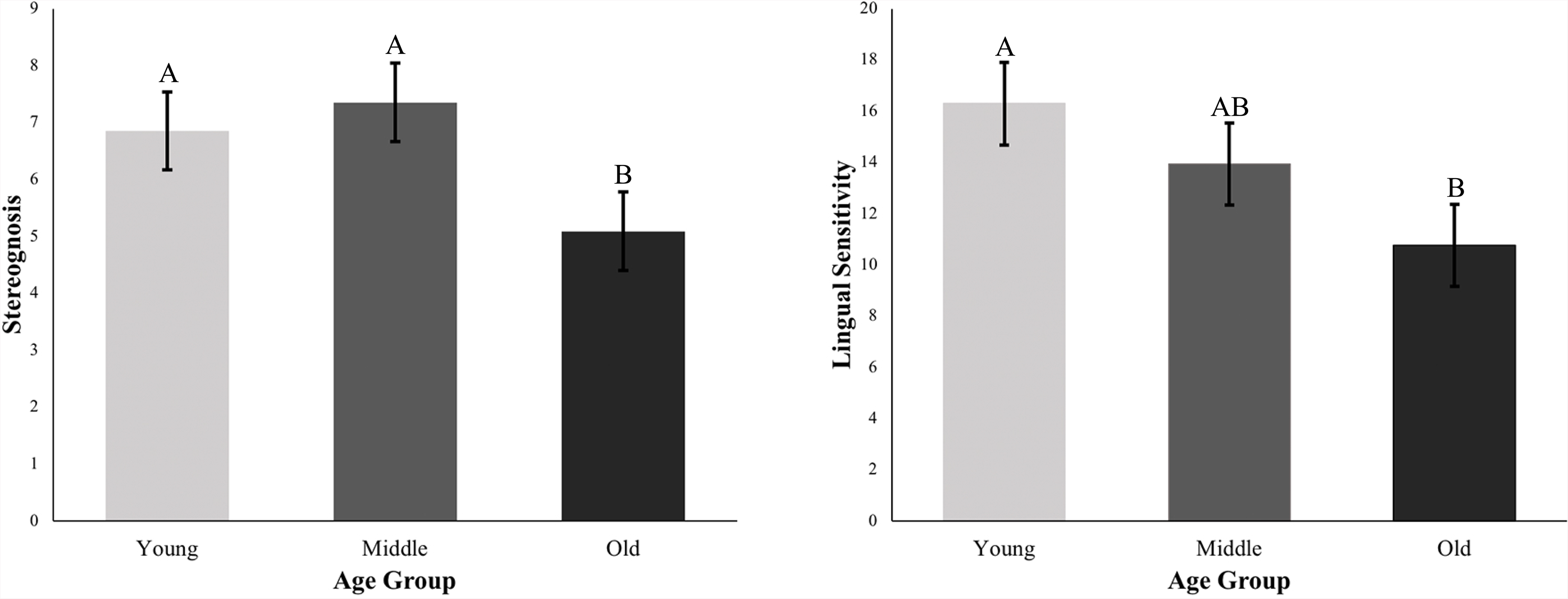
Mean values of letters and shapes correctly identified by age group, letter groupings specify significant difference (p<0.05) using Tukey’s adjustment.

Chewing efficiency was found not to differ across age groups (F_2,95_ = 1.48, p = 0.23). However, in observing the distributions of chewing efficiency by age group, it can be noted that a bimodal distribution is observed in the older age group, as shown in Figure 4.

**Figure 4.**
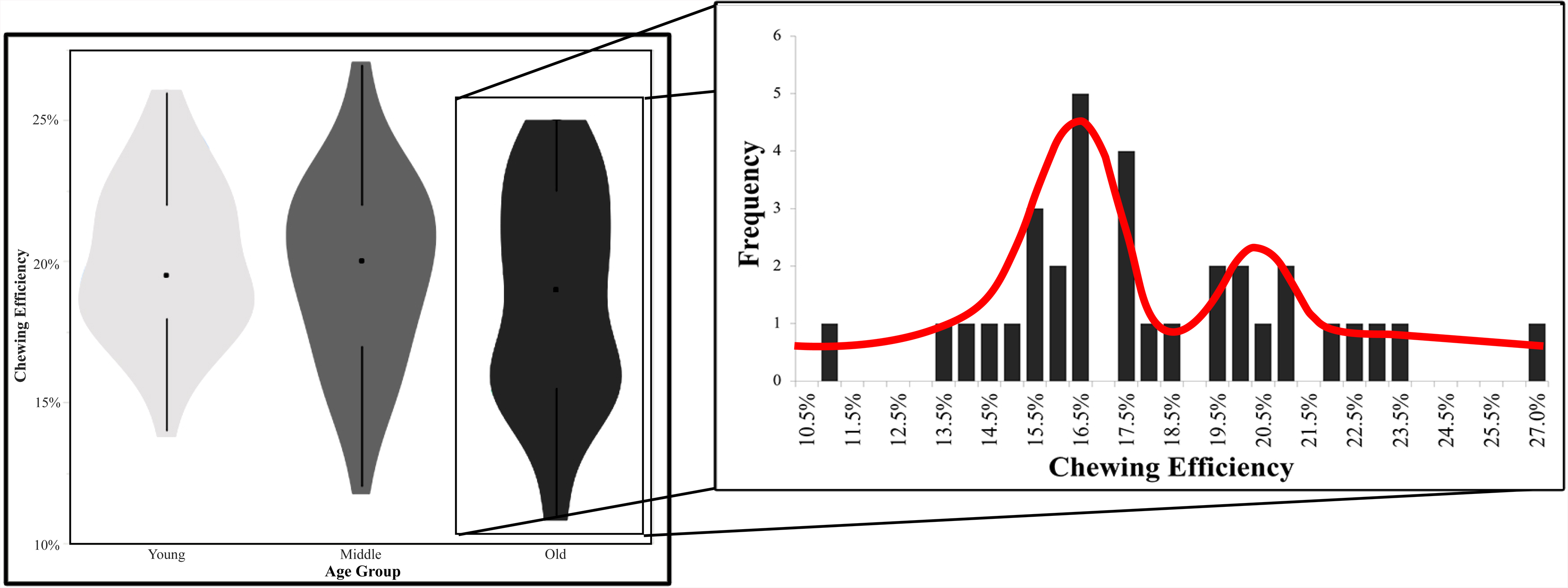
Distributions of chewing efficiency values for each of the three age groups.

### Dental Status

There was a significant effect on chewing efficiency by dental status, such that as dental status declines so does the observed chewing efficiency (r^2^ = 0.13 p=0.0268). When comparing those with notably compromised dental status (i.e. one or more missing teeth) to participants with a healthy dental status, a significantly lower chewing efficiency was found in those with missing teeth (F_1,92_ =8.59, p = 0.0043). Alternatively, with lingual sensitivity and stereognosis measures there was no significant relationships found with chewing efficiency (p=0.2396 and 0.1820). Further investigations into dental status were focused on the older adult population, since there were very few participants with notably compromised dental status in the younger age groups. Within the older adult group, it was revealed that there was no significant effect of dental status on bite force sensitivity between those with a healthy dental status and compromised participants (F_1,27_ = 4.1237, p = 0.0522). Although, it was found that chewing efficiency was significantly lower in those older adults with a compromised dental status (F_1,27_ = 5.60, p = 0.0254).

### Relationship between measurements

Increases in age were found strongly correlated with decreases in dental status (r = −0.6004, p < 0.0001). Likewise, dental status was moderately correlated with increased chewing efficiency (r = 0.2776, p = 0.0057). In looking at the specific associations of the test methods used, stereognosis showed a strong negative correlation with age (r = −0.4136, p < 0.0001) and a more moderate positive correlation with dental status (r = 0.2749, p = 0.01). Lingual sensitivity was also moderately correlated with age (r = −0.3693, p = 0.01). Both lingual sensitivity and stereognosis scores were strongly correlated with each other (r = 0.4645, p < 0.0001).

Further investigation into the different relationship that sensitivity tests were having with masticatory performance, among the older age group, bite force sensitivity (r = −0.4943, p = 0.0035) as well as dental status (r = 0.4144, p = 0.0165) were both significantly correlated with chewing efficiency. These findings highlight the multifaceted nature of oral sensitivity/processing and the need for multiple methods to comprehensively characterize oral tactile sensitivity. Among the two youngest groups, increases in bite force sensitivity was shown to significantly associate with higher chewing efficiency (r^2^ = 0.0729, p = 0.0297); and, in the older age group a significant association between chewing efficiency and bite force sensitivity was also found (r^2^ = 0.2443, p = 0.0035). However, the relationship between bite force sensitivity and chewing efficiency is in the opposite direction for the older age group (i.e. as bite force sensitivity decreases, chewing efficiency increases), therefore canceling out any effect seen across the whole participant pool (Figure 5). Total oral sensitivity relates to chewing efficiency similarly in young and middle age groups, but older adults show a different relationship.

**Figure 5.**
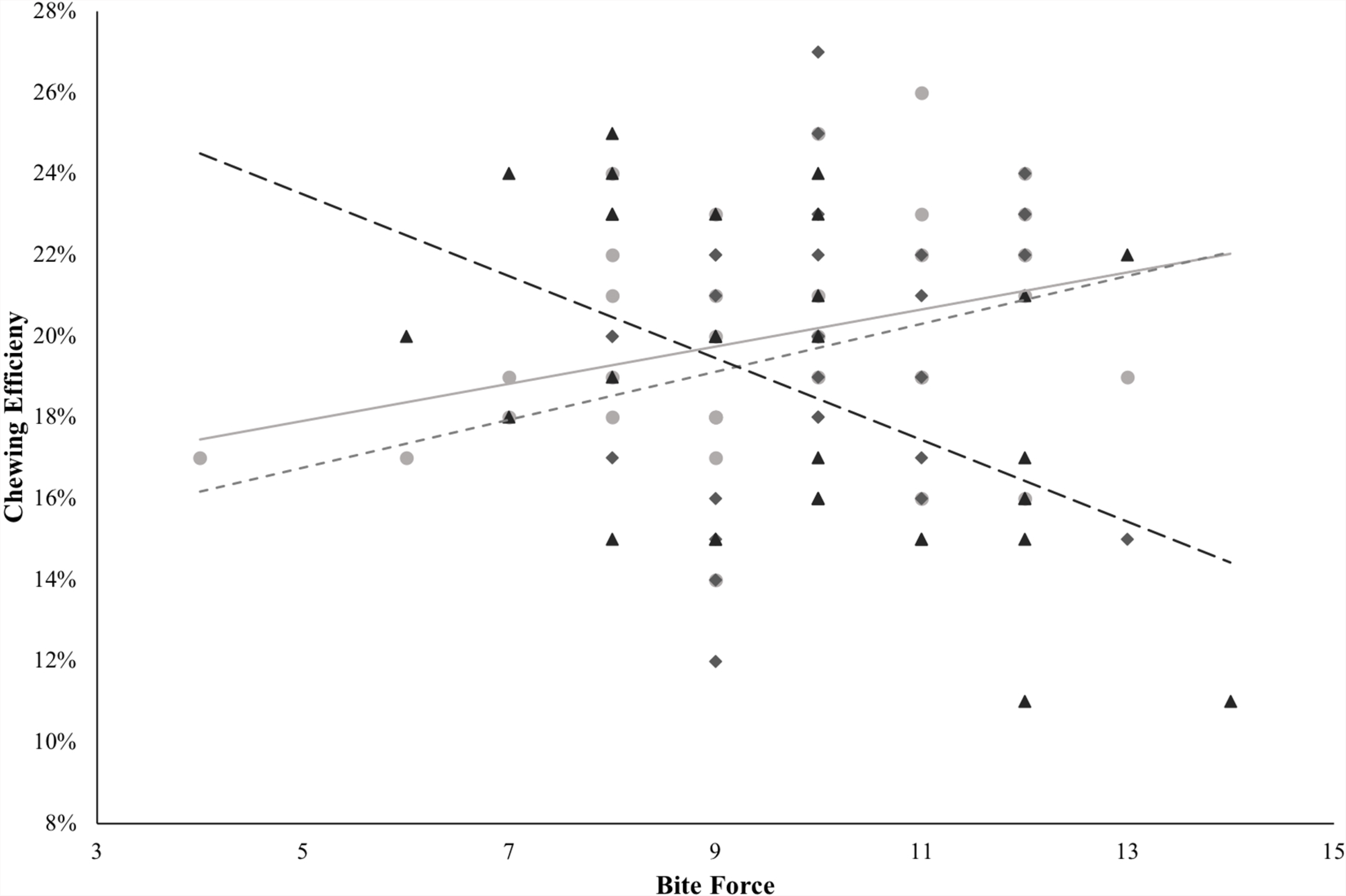
Linear Regression of chewing efficiency by pressure sensitivity, grouped by age (Young •, Middle ♦, Old ▴), showing an inverse trend as age increases.

## Discussion

The present results showed that as the population ages, there are different rates of sensitivity and proficiency decline with one group showing minimal sensory decline and another displaying notable declines. This phenomenon is also observed in other food-related sensory systems for example, olfactory sensitivity remains normal in portions of the aging population, while others exhibit a drastic loss (Murphy et al., 2002). The finding that sensitivity decrease as the population ages is in agreement with previous findings (Bangcuyo & Simons, 2017; Calhoun et al., 1992; Linne, 2017; Wada et al., 2017). Chewing efficiency of younger participants is not significantly different than those ranging from >62 years of age, showing that compensatory strategies are being utilized in the older adults. While, degradation of dental status linked to aging is not the main factor of oral sensitivity, as measured through oral lingual sensitivity testing. These findings reinforce that a host of oral sensory processing factors must be used in order to measure oral sensitivity. Even in older participants with a dental status ranging from minimal natural teeth without prosthetics to fully dentured, they performed well at pressure discrimination while they scored lower on all other tests.

Previous attempts to relate mastication ability to oral sensitivity have shown a stronger link between pressure-related measures of oral tactile sensitivity in comparison to those measures of surface sensitivity (Engelen et al., 2004). This study did not find pressure sensitivity, using bite force sensitivity, to be significantly related to mastication ability. Furthermore, bite force sensitivity is conserved as teeth are removed, pointing towards the possibility of receptors unrelated to the periodontal ligament being involved in bite force sensitivity. Although strong correlations were not found between sensitivity tests and mastication proficiency (as measured by chewing efficiency), it was noted that dental status is a significant factor in explaining the variance within chewing efficiency in the older group.

Of the three oral sensitivity tests, stereognosis was the best predictor of chewing efficiency. This indicates that the tongue proprioceptive ability, which is crucial in orienting and identifying stereognosis stimuli, may be a determining physiological factor in masticatory proficiency and further research is needed to determine the oral physiology and the masticatory feed-back loop inputs. Tongue pressure has been measured with gels of varying initial consistency using measurements of force exerted on the hard pallet and how this relates to particle size reduction, and therefore mastication, using multiple oral processes (Yokoyama et al., 2014). The tactile modalities used in this study showed a relationship with oral sensitivity and chewing efficiency, while novel techniques such as those measuring pressure sensitivity of the premandible muscle through bite failed to show a significant relationship. Bite force sensitivity was not correlated with either oral tactile measures, chewing efficiency, age, or dental status, demonstrating that bite force sensitivity measurements are likely measuring a different physiological ability from the lingual sensitivity and stereognosis measurements. These findings are in line with previous studies looking at relationships between different measures of oral sensitivity. Engelen et al. found no correlation between oral spatial acuity and oral size acuity, creating consistent evidence that oral sensitivity is multidimensional and cannot be comprehensively characterized by a single physiological assessment (2004).

This study highlights that many factors must be taken into consideration when understanding the abilities, regardless of task, of older populations. Factors such as independency level, medications, disease, nutritional deficiencies, and life style have been linked to oral perception decline (Song et al., 2016). These factors are essential at understanding perception, for example older adults have been found to have mixed responses to enhanced flavors, irritants, and modified textures that are commonly used to make products palatable for this demographic. With healthy independently living individuals having no preference or decreased liking for these modified foods, while individuals in assisted care with declining oral health prefer these modifications. In order to better meet the expectations of the desired population, work should be done to ensure the demographic is well documented and understood.

### Limitations

It was noted that even though chewing efficiency photos were taken in a controlled environment throughout the study, there were color temperature differences in the final photographs; which resulted in varying selections of blue pixels. This limitation was mediated by the use of multiple tolerances, yet room for improvement still exists. Chewing efficiency measurements were also found to be very similar, leading to possible range restriction when attempting to build relationships relating measures of oral sensitivity to chewing efficiency. Future work should be vigilant of condensed values for chewing efficiency.

The lack of a relationship between the bite force sensitivity and chewing efficiency may be due to the fact that the foam used in this study can undergo oxidation when exposed to light for prolonged periods of time, which could have resulted in a change of observed hardness over the course of the study. Oxidation of samples was mitigated by using colored containers to store samples prior to being prepped; prepped samples were used within a week, and were discarded if discoloration was observed, samples were kept in opaque containers to prevent prolonged exposure to UV light. In future studies, different types of foams could be used that have been engineered to be more resistant to oxidation from many common sources such as heat and light (Christopher, 2015).

## Conclusion

Our results show that multiple factors contribute to masticatory performance, as measured by chewing efficiency. Lingual acuity and stereognosis showed the highest correlation with chewing efficiency and appear to be the most reliable measurements, while bite force sensitivity did not show any relationship. Further research is required to quantify the relationship between physiological measures and oral sensitivity and processing ability. While some methods such as monofilaments and premandible muscle sensitivity testing have not shown promising results, modifications to these concepts may still lead to viable research. Furthermore, the tongue’s contribution to mastication ability appears to be highly correlated, showing that tongue movements or force may be a key physiological measure in future studies.

